# Mixed representations of sound and action in the auditory midbrain

**DOI:** 10.1101/2023.09.19.558449

**Authors:** GL Quass, MM Rogalla, AN Ford, PF Apostolides

**Affiliations:** Kresge Hearing Research Institute, Department of Otolaryngology – Head & Neck Surgery, University of Michigan Medical School, Ann Arbor, Michigan 48109, United States; Department of Molecular and Integrative Physiology, University of Michigan Medical School, Ann Arbor, Michigan 48109, United States

**Keywords:** Inferior Colliculus, Mixed Selectivity, Calcium Imaging, Population Analysis, Mouse

## Abstract

Linking sensory input and its consequences is a fundamental brain operation. Accordingly, neural activity of neo-cortical and limbic systems often reflects dynamic combinations of sensory and behaviorally relevant variables, and these “mixed representations” are suggested to be important for perception, learning, and plasticity. However, the extent to which such integrative computations might occur in brain regions upstream of the forebrain is less clear. Here, we conduct cellular-resolution 2-photon Ca^2+^ imaging in the superficial “shell” layers of the inferior colliculus (IC), as head-fixed mice of either sex perform a reward-based psychometric auditory task. We find that the activity of individual shell IC neurons jointly reflects auditory cues and mice’s actions, such that trajectories of neural population activity diverge depending on mice’s behavioral choice. Consequently, simple classifier models trained on shell IC neuron activity can predict trial-by-trial outcomes, even when training data are restricted to neural activity occurring prior to mice’s instrumental actions. Thus in behaving animals, auditory midbrain neurons transmit a population code that reflects a joint representation of sound and action.

**Significance Statement:** Neurons in IC’s superficial “shell” layers preferentially project to higher-order thalamic nuclei that are strongly activated by sounds and their behavioral consequences. This integrative computation is thought critical for a variety of behaviorally relevant functions, such as establishing learned sound valence. However, whether such “mixed representations” reflect unique properties of thalamocortical networks, or rather are inherited from afferent inputs, is unclear. We show that in behaving mice, many shell IC neurons are modulated by sounds and mice’s actions. Consequently, shell IC population activity suffices to predict behavioral outcomes even prior to the goal-directed action. Our data thus establish shell IC nuclei as a novel, ascending source of mixed representations for the thalamocortical system.

## Introduction

Choosing the appropriate behavioral response to appetitive or aversive stimuli confers a survival advantage. To achieve this, neural circuits must be capable of linking external sensations, instrumental actions, and their behaviorally relevant consequences. One solution is for distinct sensory and behaviorally relevant pathways to converge upon a common target region, thereby enabling postsynaptic ensembles to jointly encode sensations and their consequences such as reward, punishment, or goal-directed actions. Indeed, such “mixed-selectivity” to sensory and behavioral variables is well-documented in the thalamus (L. Chen et al., 2019; Gilad et al., 2020; Hu, 2003; Jaramillo et al., 2014; Komura et al., 2001; Ryugo & Weinberger, 1978) and neo-cortex, and might contribute to the computational power of these high-level circuits (Naud & Sprekeler, 2018; Parker et al., 2020; Rigotti et al., 2013; Saxena et al., 2022; Stringer et al., 2019). However, whether such joint representations reflect unique integrative computations of the thalamo-cortical system, or are inherited from afferent inputs, is unknown.

The inferior colliculus (IC) is a midbrain hub that transmits most auditory signals to the forebrain (Aitkin et al., 1981; Aitkin & Phillips, 1984; LeDoux et al., 1985, 1987; Coleman & Clerici, 1987). It is sub-divided into primary central and surrounding dorso-medial and lateral “shell” nuclei whose neurons preferentially project to primary and higher-order medial geniculate body (MGB) of the thalamus, respectively (C. Chen et al., 2018; Mellott et al., 2014; Winer et al., 2002). Interestingly, lesions to the shell IC or their afferent inputs apparently do not cause central deafness, but rather seemingly impair certain forms of learned auditory associations (Jane et al., 1965; Bajo et al., 2010). In tandem with their anatomical connectivity to non-lemniscal thalamic regions, these results suggest that shell IC neurons may be involved in higher-order auditory processing and learned sound valence.

Accordingly, shell IC neuron activity is modulated by behavioral engagement, movement, and reward expectation (C. Chen & Song, 2019; Lee et al., 2023; Shaheen et al., 2021; van den Berg, 2021, De Franceschi & Barkat, 2021). Although some of these effects can be explained by an arousal-mediated scaling of acoustic responses (Joshi et al., 2016; Saderi et al., 2021), whether the shell IC additionally transmits high-level signals related to behavioral outcome and goal-directed actions is less clear. Interestingly, higher-order MGB neurons jointly encode combined sound and behavioral outcome signals, which may serve important learning related functions (Edeline & Weinberger, 1992; McEchron et al., 1995; Mogenson et al., 1980; Ryugo & Weinberger, 1978; Schultz et al., 2003; Taylor et al., 2021). However, whether such integrated representations of acoustic and behaviorally relevant information are already present in upstream shell IC neurons is unknown.

Here we use 2-photon Ca^2+^ imaging to record shell IC neurons as head-fixed mice engage in an appetitive auditory, Go/NoGo task. We find that shell IC populations encode sound- and behaviorally relevant information that is predictive of mice’s instrumental choice on a trial-to-trial basis. Thus, the auditory midbrain broadcasts a powerful mixed representation of sound and outcome signals to the thalamocortical system.

## Materials And Methods

### Animal subjects and handling

All procedures were approved by the University of Michigan’s Institutional Animal Care and Use Committee and conducted in accordance with the NIH’s guide for the care and use of laboratory animals and the Declaration of Helsinki. Adult CBA/CaJ x C-57Bl-6/J mice were used in this study (n=11, 5 females, 70-84 days postnatal at time of surgery). These hybrids do not share the Cdh23-mutation that results in early-onset presbycusis in regular C57Bl-6 mice (Frisina et al., 2011; Johnson et al., 1997; Kane et al., 2012). Following surgery, mice were single-housed to control water-deprivation and to avoid damage to surgical implants. Cages were enriched (running wheels, nest building material), kept in a temperature-controlled environment (24.4°C, 38.5% humidity) under an inverted light-dark cycle (12 h/12 h), and mice had olfactory and visual contact to neighboring cages. Three animals entered the experiment after having spent three prior sessions where they were passively exposed to different sound stimuli than employed in the current study (Shi et al., 2023).

### Surgery

Mice were anesthetized in an induction chamber with 5% isoflurane vaporized in O_2_, transferred onto a stereotaxic frame (M1430, Kopf Instruments, Tujunga, CA, USA), and injected with carprofen as a pre-surgical analgesic (Rimadyl, Parsippany-Troy Hills Township, NJ, USA; 5 mg/kg s.c.). During surgery, mice were maintained under deep anesthesia via continuous volatile administration of 1-2% isoflurane. Body temperature was kept near 37.0°C via a closed loop heating system (M55 Harvard Apparatus, Holliston, MA, USA), and anesthesia was periodically confirmed by absence of leg-withdrawal reflex upon toe pinch. The skin above the parietal skull was removed, and a local anesthetic was applied (Lidocaine HCl, Akorn, Lake Forest, IL, USA). The skull was balanced by leveling the vertical difference between Lambda and Bregma coordinates, and a 2.25-2.5 mm diameter circular craniotomy was carefully drilled above the left IC at Lambda −900 μm (AP) / −1000 μm (LM). The skull overlying the IC was removed, and pAAV.Syn.GCaMP6f.WPRE.SV40 (AAV1, titer order of magnitude 10^-12^ Addgene) was injected 200 μm below the dura at 4 different sites (25 nL each; 100 nL total) across the medial lateral axis of the IC using an automated injection system (Nanoject III, Drummond, Broomall, PA, USA). In three cases, pAAV.syn.jGCaMP8s.WPRE (AAV1, titer order of magnitude 10^-12^, Addgene) was injected. A custom-made cranial window insert, consisting of three circular 2 mm glass coverslips stacked and affixed to a 4 mm diameter glass outer window, was then inserted in the craniotomy. The cranial window was affixed to the skull, sealed with cyanoacrylate glue (Loctite, Westlake, OH, USA) and a titanium head bar was mounted on the skull with dental cement (Ortho-Jet, Wheeling, IL, USA). Animals received a post-surgical subcutaneous injection of Buprenorphine (0.03 mg/kg s.c., Par Pharmaceuticals, Chestnut Ridge, NY, USA). Mice received Carprofen injections (5 mg/kg, s.c., Spring Meds, Sioux Falls, SD, USA) 24 and 48 hours following surgery.

### Behavior protocol

After a minimum of 14 days recovery from surgery, mice were water restricted (1-1.5 ml/day) and maintained at >75 % initial body weight. Mice were habituated to the experimenter, the experimental chamber, and the head-fixation. During the habituation- and experimental sessions, mice sat in an acrylic glass tube in a dark, acoustically shielded chamber with their heads exposed and fixed, and a lick spout in comfortable reach. Following 7 days of water restriction and acclimation, mice were trained daily in a reward-based, operant Go/NoGo paradigm (Figure 1A, B), controlled by a Bpod state machine (Sanworks, Rochester, NY, USA) run with Matlab (version 2016b, MathWorks, Natick, MA, USA). Sounds were generated in Matlab at a sampling frequency of 100 kHz and played back via the Bpod output module. A sound was presented from a speaker (XT25SC90-04, Peerless by Tymphany, San Rafael, CA, USA) positioned 30 cm away from the animal’s right ear (1 s duration, 70 dB SPL, calibrated using a 1/4” pressure-field microphone [Bruel & Kjaer, Nærum, DK]). Licking behavior was recorded for the entire trial time using a light-gate in front of the spout, and sampled down to 7.3 Hz (C57BL/6J lick frequency, Boughter et al., 2007) and binarized offline. During Go-trials, licking a waterspout during a 1 s “answer period” following sound offset resulted in delivery of a reward (10 % sucrose-water droplet gated through a solenoid valve). During NoGo-trials, mice had to withhold licking during the answer period and false alarms were punished with an increased inter-trial interval (“timeout”). Licking at any other point in the trial had no consequence. Thus, inter-trial intervals were 13-15 s following all Go- and correctly answered NoGo-trials, and 18-20 s for incorrectly answered NoGo-trials. Inter-trial intervals were kept this long to avoid photo-bleaching and laser damage to the tissue, while approximately balancing laser-on-time (8 s) and laser-off-time (5-12 s).

**Figure 1.**
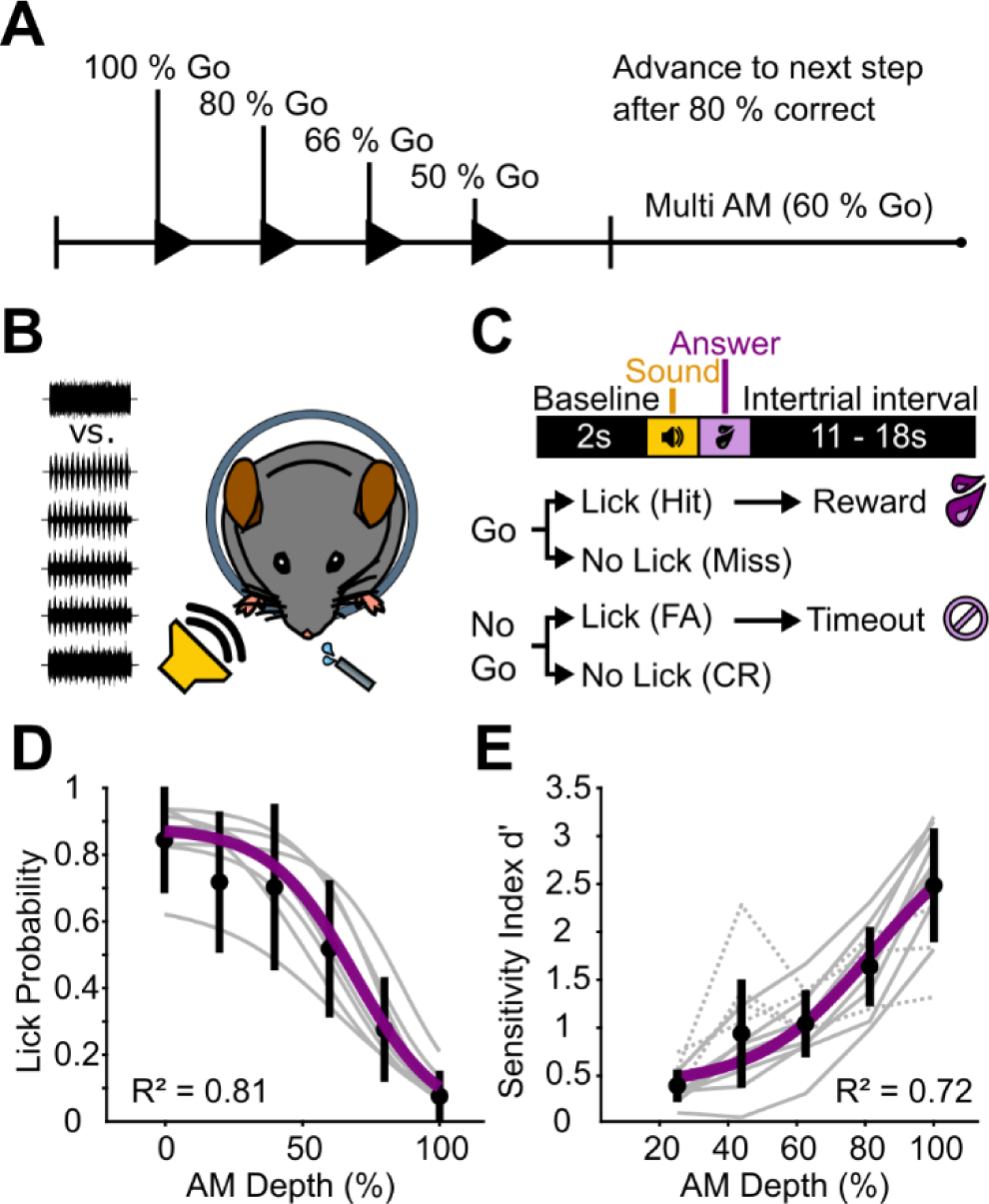
Mice discriminate sAM-noise from unmodulated noise in a modulation depth-dependent manner. A – Experiment structure. Head-fixed mice were trained to discriminate between 0 % and 100 % sAM depth. We progressively reduced the ratio of Go to NoGo-trials as mice’s task performance increased. B –Upon reaching criterion (see Results text), mice engage in a “multi-sAM depth” version of the task where the modulation depth of the NoGo sound was varied on a trial-by trial basis C – Trial structure. After a 2 s baseline, a sound was presented for 1 s. Licking a waterspout during a 1 s answer period following sound offset was rewarded with a drop of sugar water on Go-trials, and punished with a 5 s timeout on NoGo-trials. Licking at any other point during the trial had no consequence. D – Fitted lick probability during the answer period as a function of sAM depth for all mice during multi-sAM sessions. Gray lines are individual animals, black circles and lines are mean ± standard deviation of each sAM depth, purple line is the mean fit. E – d’ per sAM depth for all mice (Gray lines). Solid and dashed lines are mice that received unmodulated noise and sAM noise as Go-stimuli, respectively. Black circles and lines are mean ± standard deviation for each sAM depth, purple line is the mean fit.

All mice were trained according to the same protocol: In the first stage, only Go-stimuli were presented, and rewards were manually triggered by the experimenter, so that mice learned to associate the Go-stimulus with a water reward (shaping, usually continuously for the first 10 trials, followed by slowly decreasing manual reward delivery until trial 50 during initial Go-only sessions). This procedure was repeated over multiple sessions until an association was present, determined by reaching a criterion of 80 % response rate without shaping in 2 consecutive sessions. Next, the NoGo-stimulus was introduced. In this stage, the number of NoGo presentations was gradually increased from 20 %, to 33 %, to 50 % if mice responded correctly on 80 % of trials during a session. A typical session contained around 200 trials and lasted for up to 1 h. During these training stages, (sAM = NoGo, n = 7)the Go stimulus was a broad-band noise burst (BBN, 4 – 16 kHz), and the NoGo stimulus was an amplitude-modulated BBN modulated at 100 % depth and a modulation frequency of 15 Hz (sAM, 4 - 16 kHz BBN carrier). To ensure that mice attend to the temporal envelope modulation of the stimulus, we trained a subgroup of 4 mice on reversed stimuli (sAM = Go). The data were analyzed jointly unless otherwise noted.

After reaching 80% correct in the 50/50 Go/NoGo stage for 2 consecutive sessions, we varied the amplitude modulation depth (NoGo-stimulus for group 1, Go-stimulus for group 2) from 20% to 100% in 20% steps. Mice performed 6 - 7 sessions in this paradigm, with a typical session containing around 350 trials and lasting for up to 1.5 h. If an animal that had learned the task produced misses for 6 Go-trials in a row, the session was terminated since these trials were indicative of a lack of motivation and licking. Due to the pseudo-randomized trial order, this criterion was reached over a maximum of 14 consecutive trials once a mouse stopped licking. Thus, the final 14 trials of each session were discarded from all analyses.

Water intake during the task was estimated by measuring the mice’s weight difference (including droppings) before and after each session. Mice received supplementary water if they consumed less than 1 ml during the session. Upon conclusion of the experiment, mice received water *ad libitum* for at least two days, were deeply anesthetized via an overdose of isoflurane, and transcardially perfused with formalin.

### Behavior analysis

Lick responses to assign trial outcomes were counted only during the reward period (1 s after sound offset). Licking at any point during the reward period during Go-trials resulted in a Hit and was immediately rewarded. Not licking during this period was scored as a Miss. Licking during the reward period during NoGo-trials was counted as a False Alarm (FA) and resulted in a timeout, and not licking was scored as a Correct Rejection (CR) and was not rewarded nor punished. Licking at any other point during any trial had no consequence.

The sensitivity index d’ was calculated as d’ = z(hit rate) - z(false alarm rate), where z(hit rate) and z(false alarm rate) are the z-transformations of the hit rate and the false alarm rate, respectively. Global lick rates pooled from all sessions were fitted per animal with a 4-parameter logistic equation (sigmoid fit), and the perceptual threshold was defined as the modulation depth at which half-maximal lick probability was reached.

### Ca^2+^ imaging

Movies were acquired at a frame rate of 30 Hz (512 x 512 pixels) using a resonance-scanning, 2-photon microscope (Janelia Research Campus’ MIMMs design; Sutter Instruments, Novato, CA, USA) equipped with a 16x water immersion objective (Nikon, 0.8 NA, 3 mm working distance) and a GaAsP photomultiplier tube (Hamamatsu Photonics, Hamamatsu, Japan). The microscope was located in a custom-built, sound- and light-attenuated chamber on a floating air table. GCaMP6f or −8s were excited at 920 nm using a Titanium-Sapphire laser (30 – 60 mW absolute peak power, Chameleon Ultra 2, Coherent, Santa Clara, CA, USA). Images were acquired for 8 s per trial from the same field of view in each session (determined by eye using anatomical landmarks), with a variable inter-trial-interval (see Behavior protocol). Recording depth from dura was variable between animals and chosen by image quality and number and responsiveness of neurons (tested live), but generally kept between 20 and 55 µm. Behavioral data (licks) were recorded simultaneously through Matlab-based wavesurfer software (Janelia Research Campus) and synchronized with the imaging data offline.

### Ca^2+^ imaging analysis

We used the Python version of Suite2p to motion-correct the movies, generate regions of interest (ROIs), and extract fluorescence time series (Pachitariu et al., 2016). ROIs were manually curated by the experimenter to exclude neurites without somata, and overlapping ROIs were discarded if they could not be clearly separated. Raw fluorescence time series were converted to ΔF/F by dividing the fluorescence by the mean fluorescence intensity during the 2 s baseline period on each trial, subtracting the surrounding neuropil signal scaled by a factor of 0.7, and smoothing the traces using a 5-frame gaussian kernel. ΔF/F traces and behavioral data were then analyzed using custom Matlab routines (available upon request). To determine significantly responding ROIs, we used a bootstrapping procedure based on the ΔF/F “autocorrelation” across similar trial types (Geis et al., 2011; Wong & Borst, 2019). Briefly, the average correlation over either the sound- or the answer period of each matching pair of trials with the same stimulus was compared to its correlation with a randomly sampled signal from the same trials 10000 times. The p-values were then computed as the fraction of these randomly sampled signals with greater correlation than the real data, and corrected for multiple comparisons using the Bonferroni-Holm method. Importantly, this method measures trial-to-trial consistency, and not response onset or strength. Thus, prolonged, but consistent sound responses may occasionally lead to significantly outcome-responsive neurons. Since decreases in fluorescence can be difficult to interpret specifically for tuning analyses, we used t-SNE (van der Maaten & Hinton, 2008) and k-means clustering (2 clusters) in the tuning analyses to separate sound-excited from sound-inhibited neurons by their average ΔF/F waveform, and only analyzed sound-excited neurons. In population analyses (PCA and SVM), all neurons were used, regardless of whether they were significantly responding, sound-excited, or sound-inhibited, according to our analyses. The outcome selectivity index of each neuron was calculated as follows: We first averaged ΔF/F traces of Hit, Miss, CR, and FA trials. We then measured the absolute value integrals of each average waveform from sound onset to 1 s after the answer period. Outcome selectivity indices for Go and NoGo trials were calculated as (Hit – Miss)/(Hit + Miss) and (CR – FA)/(CR + FA), respectively.

### Lifetime sparseness

As an additional measure for neuronal selectivity, we computed the lifetime sparseness per neuron, which describes a neuron’s general activity variance in response to an arbitrary number of stimuli (Vinje & Gallant, 2000). Here, we computed the lifetime sparseness separately for modulation depth and trial outcome:

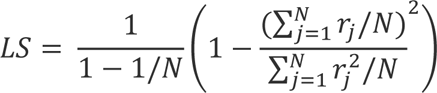

where *N* is the number of different stimuli and *r_j_* is the mean peak ΔF/F response to stimulus *j* from sound onset to +2 s. Before computing the lifetime sparseness, we set all negative ΔF/F traces (neurons that reduced their firing relative to baseline) to 0 to keep the lifetime sparseness between 0 and 1. Lifetime sparseness is 0 when a neuron responds to all stimuli with the same peak ΔF/F response, and 1 when it only responds to a single stimulus.

### Support vector machine classifier

The support vector machine (SVM) classifier was generated in Matlab using the classification learner app, with the “templateSVM” and “fitsvm” or “fitecoc” functions as the skeleton for binary and multi-label classification, respectively . In all cases, we used a linear kernel and the sequential minimal optimization algorithm to build the classifier. We used the ROIs as individual predictors and one of several sets of variables as classes: behavioral outcome (hit, miss, false alarm, correct rejection), action during answer period (lick, no lick), stimulus identity (AM depth), stimulus category (Go-stimulus, NoGo-stimulus), and presence of error (correct response, incorrect response), using equal priors. The integral of the ΔF/F traces over 100 ms was used as the input data, and the classifier was constructed and trained on each individual session using an equal number of Go- and NoGo-trials. We used periods of 100 ms in steps of 100 ms over the whole signal to extract the information content in the signal at each period. Thus, at each time point t, the classifier has access to the integral of the activity from t to t + 100 ms. For the “First Lick Accuracy”, the 100 ms preceding the first lick that occurred at least 100 ms after stimulus onset was used. If no lick was present during a trial, the median first lick time of all licked trials of that session was used instead. We used 5-fold validation to determine the decoding accuracy per session, i.e., 5 randomly sampled portions of 80 % of trials as training data, and the remaining 5 times 20 % as test data. The accuracy is then given as the mean decoding accuracy among those five folds. Because the number of trials per class is not always balanced, we computed the “Balanced Accuracy”, which is calculated differently for binary (lick/ no lick, Go/ NoGo, error/ no error) and nonbinary problems (trial outcome, AM depth). For binary problems, the balanced accuracy is defined as the number of true positives plus the number of true negatives, divided by 2. Thus, the balanced accuracy normalizes the accuracy to 50 % at chance level. For non-binary problems, it is defined as the mean of the micro-recalls (recall/class, see below), and chance level is 1 divided by the number of classes. All data is presented as “Balanced Accuracy”.

As controls, we computed the “Shuffled” and “Shuffled Balanced” Accuracies, where the trials and the class labels are shuffled prior to classifier training. This method thus reflects a real chance level. To prevent overfitting to individual, strongly selective neurons, we included a “dropout”-rate of 10 % by setting the ΔF/F traces of 10 % randomly sampled ROIs in each trial to 0 during the training.

We further assessed the quality of our classifiers by computing the weighted precision (positive predictive value, or exactness, the number of true positives divided by the number of all positives, weighted by class prevalence) and weighted recall (sensitivity or completeness, the number of true positives divided by the number of true positives and false negatives, weighted by class prevalence), and then computing the weighted F1-score (the harmonic mean of the two, Hand et al., 2001; Rijsbergen, 1979) and the AUC as the mean of the AUCs per class (area under the receiver-operating-characteristic, Huang et al., 2003). This information was used to compute the balanced accuracy for multiclass problems.

### Principal component analysis

We performed a population principal component analysis (PCA) using individual ROIs as observations, and the trial-averaged ΔF/F samples as individual variables using Matlab’s “pca”-function with the default parameters. To compare the differences in the multidimensional neural trajectories, we computed the mean weighted Euclidean distances (_w_euΔ) of hit- and miss-, and correct rejection- and false alarm-trials, respectively. The _w_euΔ was obtained by computing the Euclidean distance between each component of the neural trajectories at each point in time, weighted by the amount of variance explained by each component, resulting in a weighted distance-vector per component. These were then summed up to a single ∑_w_euΔ-curve per session that is proportionate to the general difference in network activity and normalized to the intra-session variance.

### Statistics

All statistical analyses were run in Matlab. Significance levels *, **, and *** correspond to p-values lower than 0.05, 0.01, and 0.001, respectively. Data were tested for normality using the Kolmogorov-Smirnov test and nonparametric tests were used when the data were not normally distributed. All descriptive values are mean and standard deviation unless otherwise noted. P-values were corrected for multiple comparisons where appropriate using the Bonferroni-Holm method. Sample sizes were not pre-determined.

## Results

### Head-fixed mice discriminate amplitude-modulated from unmodulated noise in an operant task

Water-deprived, head-fixed mice (N = 8) were trained to discriminate the presence or absence of 15 Hz sinusoidal amplitude modulation (sAM) in a 1 s band-limited noise (4 – 16 kHz, 70 dB SPL) using an operant Go/NoGo paradigm (see Materials & Methods for full description of training regimen). On Go-trials, the noise carrier sound was presented without amplitude modulation (0 % sAM depth). Licking a waterspout within a 1 s “answer period” following sound offset was scored as a “Hit” outcome, and rewarded with a drop of 10 % sucrose water. Withholding licking during the answer period of Go trials was scored as a “Miss” outcome and neither punished nor rewarded. On NoGo-trials, the noise carrier was fully amplitude-modulated (100 % sAM depth), and mice had to withhold licking during the answer period; these “correct rejection” outcomes were not rewarded. Licking during the answer period of NoGo trials was scored as a “false alarm” outcome and punished with a 5 s “time out” (increased inter-trial interval; Figure 1A, B, C). Mice reached a criterion expert performance of ≥80 % correctly responded trials after 13.6 ± 2.6 training sessions.

After mice reached expert performance (<20 % false alarms for two consecutive sessions), we varied the modulation depth of the sAM sound in subsequent sessions from 20 % to 100 % in steps of 20 %. False alarm rates increased in this “multi-sAM” paradigm compared to the final two sessions with only 0 and 100 % sAM depths (0.46 ± 0.16 vs 0.18 ± 0.11 respectively, mean and standard deviation), as expected from an increased perceptual ambiguity of NoGo sounds on low sAM depth trials (Figure 1D). However, mice’s Hit- and False Alarm rates remained stable across consecutive daily sessions, indicating that discriminative performance did not increase with further training on the multi-sAM paradigm (ANOVA, F(6,58) = 1.46 for factor Session #, p = 0.2065). Expectedly, false alarm rates were not evenly distributed across NoGo conditions of varying depths, and mice were more likely to lick on low sAM depth NoGo trials (mean fit half-maximal lick probability: 69 % sAM depth, Figure 1E). Because false alarm rates increased with the perceptual similarity of Go and NoGo sounds, these data argue that performance reflects mice’s attending to temporal envelope modulation.

As a separate test of whether mice were indeed attending to the discriminative sound’s temporal envelope, we trained n = 3 mice on the opposite contingency with the presence of sAM serving as the Go stimulus. These mice’s operant responses in the multi-sAM paradigm also varied in a manner expected from temporal envelope detection. To quantify the performance for all mice regardless of training contingency, we calculated the sensitivity index (d-prime) per AM depth rather than the licking probability (Figure 1F). On average, d’ steadily rose with increasing AM depth (i.e., increasing perceptual distance from the 0 % band-limited noise carrier). Because both groups appeared to use the same strategy to solve the task, we pooled the data for all outcome-related measures. Thus, head-fixed mice rapidly learn, and stably execute our sAM detection task.

### Shell IC neuron activity shows both auditory- and non-auditory selectivity

We next investigated the extent to which behaviorally relevant activity is present in the shell IC of actively listening mice. To this end, we used a viral approach to broadly express a genetically encoded Ca^2+^ indicator (GCaMP6f or −8s) in the IC, and conducted 2-photon Ca^2+^ imaging to record shell IC neuron activity as head-fixed mice engaged in the multi-sAM task (Figure 2A, B). The multi-sAM paradigm enables comparing neural activity on trials with similar sounds, but distinct trial outcomes (i.e., hits vs. misses; correct rejections vs. false alarms), thereby testing how shell IC activity varies depending on mice’s instrumental choice in the answer period.

**Figure 2.**
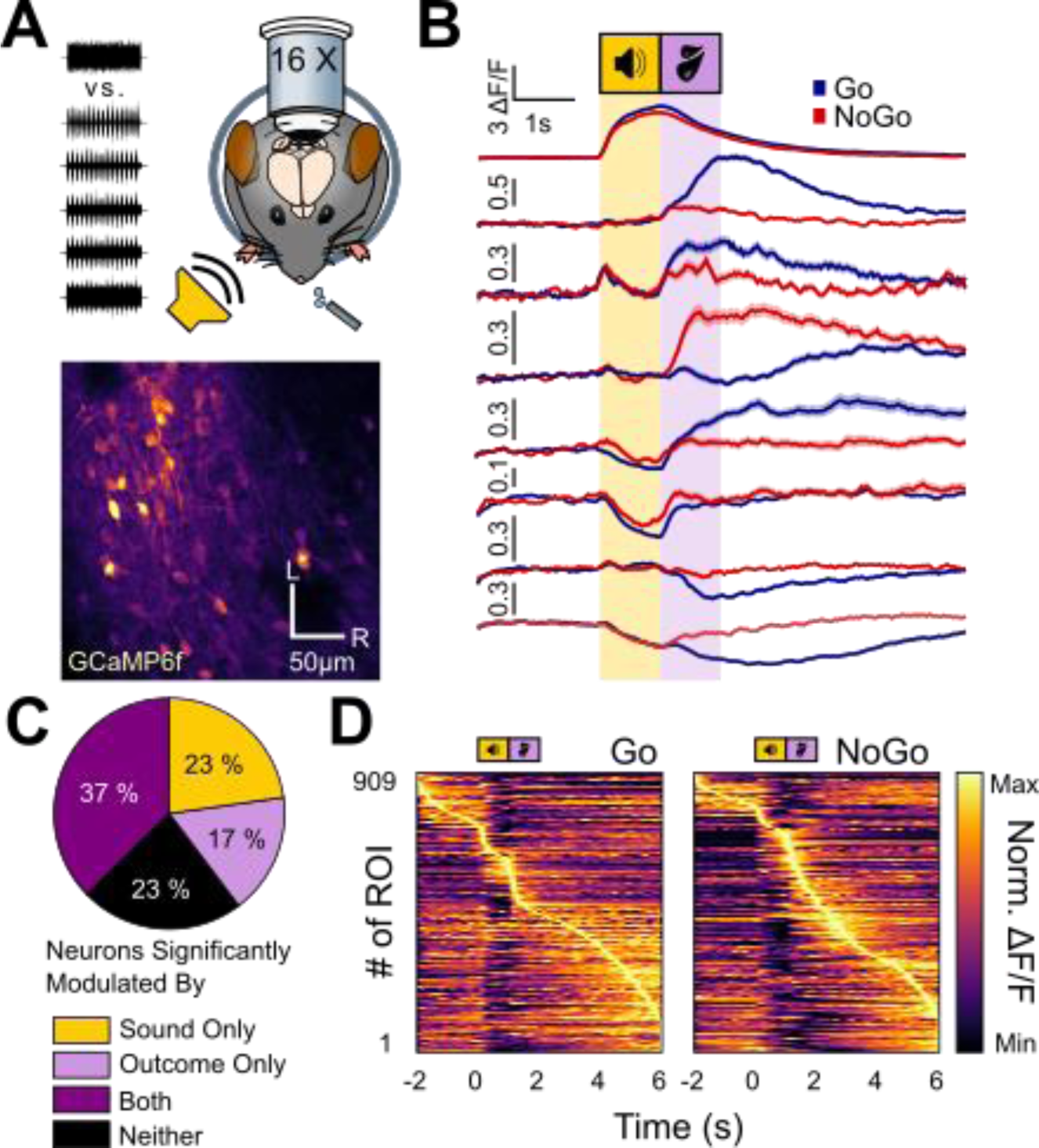
Shell IC neurons are active across the entirety of Go and NoGo trials. A – Upper panel: Experimental approach: multiphoton Ca2+ imaging was conducted in the superficial shell IC layers to record neural activity as mice engaged in the multi-sAM task. Lower panel: Example field of view from a typical session (L – lateral, R – rostral). B – Example average fluorescence traces of eight separate ROIs on Go (blue) and NoGo (red) trials. All ROIs were recorded simultaneously in the same FOV. Of note is that differential neural activity on Go and NoGo trials spans across the entire trial epoch and is expressed as both increases and decreases in fluorescence. C – The proportion of cells significantly modulated by any sound or outcome, any combination of sound and outcome, or none of those three options (n = 909). D – Distribution of activity maxima for all recorded ROIs during all mice’s first sessions sorted by trial type. The heatmaps show the average trace per ROI. Of note, most ROIs have their activity maxima after the sound termination.

We recorded n = 909 regions of interest (ROI) in n = 11 sessions from N = 11 mice (83 ± 27 ROIs per field of view, Figure 2B). We restricted all individual cell analyses to mice’s first multi-sAM session to prevent repeated measurements from the same neurons over multiple sessions. As a first pass to determine how shell IC neurons respond to task-relevant variables, we averaged each neuron’s baseline-normalized fluorescence traces (ΔF/F) separately for all Go and NoGo trials in a given session. As expected from prior imaging studies in anesthetized and passively listening mice (Barnstedt et al., 2015; Ito et al., 2014; Wong & Borst, 2019), some shell IC neurons showed strong sound-evoked fluorescence increases (Figure 2B, ROI I) or decreases (Figure 2B, ROI VI) which reflect bidirectional changes in firing rates (Wong & Borst, 2019). However, many neurons showed substantial non-auditory activity, such that maximal activity modulation occurred during the answer period following sound termination. This activity was driven by fluorescence increases (Figure 2B, ROIS II-V) as well as decreases (Figure 2B, ROIs VII and VIII), and neurons had varying degrees of selectivity for Go or NoGo trials (Figure 2B; compare ROIs II, III and IV). Shell IC neuron activity is thus bi-directionally modulated across the entire duration of behavioral trials of our task, thereby transmitting signals beyond discriminative sound cue features. Moreover many neurons showed fluorescence changes during both sound and answer periods (Figure 2B, ROIs III, V, and VIII), implying a joint coding, or mixed selectivity, for acoustic and higher-order information in shell IC neurons.

We summarized task-relevant activity by quantifying the relative proportion of sound- and answer period responsive shell IC neurons across our recordings. To this end, we employed an “autocorrelation”-bootstrapping significance analysis (Geis et al., 2011; Wong & Borst, 2019, Figure 2C) testing neuronal selectivity for sound- and answer periods using trial-to-trial correlation. This analysis suggested four major response classes of shell IC neurons: 23 % (207/909) of all recorded neurons were strictly sound responsive, 17 % (154/909) of neurons were exclusively answer period-responsive, 37 % (340/909) showed consistent sound and trial outcome responses, and 23 % (208/909) of neurons responded neither to sound nor outcome in a systematic manner detected by these analyses. Consequently, the majority of neurons in our datasets (77 %; 701/909) showed reliable activity modulation across the behavioral trials. Interestingly however, purely sound responsive neurons were a surprising minority of shell IC neurons; most neurons instead reached their activity peak after sound offset, suggesting that answer period activity is the dominant efferent signal from shell IC neuron populations under our conditions (Figure 2D).

### Sound responsive shell IC neurons are broadly tuned to sAM depth

Sound evoked spike rates of central IC neurons generally increase monotonically with higher sAM depths (Joris et al., 2004; Rees & Møller, 1983). However, non-monotonic sAM depth coding has also been reported (Preuß & Müller-Preuss, 1990), whereby neurons selectively respond to a “preferred” sAM depth akin to the non-monotonic intensity selectivity of brainstem or auditory cortex neurons (Sadagopan & Wang, 2008; Young & Brownell, 1976). We thus wondered how the shell IC neurons in our recordings encode sAM depth. To this end, we used t-SNE/k-means clustering to identify neurons showing a sound-evoked fluorescence increase (272/464), given the interpretive difficulty of fluorescence decreases (Vanwalleghem et al., 2021) and the broad selectivity of sound-evoked inhibition in shell IC neurons (Shi et al., 2023). As a first pass, we used the trial-to-trial-correlation approach to determine whether one or multiple sAM depths drove consistent responses during sound presentation. 38 % (104/272) of neurons were significantly responsive to all sAM depths (**Figure 3A**), suggesting that more than a third of sound-responsive shell IC neurons indiscriminately transmit acoustic information under our conditions. 22 % (60/272) of cells preferentially responded to low AM depths (0 % and 20 %), but not to higher ones (**Figure 3B**), and only 6 % (17/272) responded to high sAM depths (80 % and 100 %), but not to lower ones (**Figure 3C**). Only a single cell was preferentially responsive to medium sAM depths (40 % and 60 %, **Figure 3D**). The categorically broad sAM depth selectivity was also reflected in the magnitude of shell IC neurons’ responses during sound presentation, i.e., the average peak of ΔF/F traces during the sound presentation Indeed, there was no significant correlation between ΔF/F peak and sAM depth (Pearson’s ρ = 0.26, R² = 0.07, Figure 3E), suggesting that the average population activity of sound excited shell IC neurons does not systematically increase or decrease with sAM depth. Finally, we computed the lifetime sparseness of all neurons as a separate measure of tuning sharpness across all stimuli (Vinje & Gallant, 2000, Figure 3F). The population distribution of lifetime sparseness values was broad, with a low median value of 0.28 (median absolute derivation 0.18), in further agreement with weak selectivity to sAM depth. Altogether our analyses find scant evidence for non-monotonic sAM depth encoding, and furthermore indicate that most shell IC neurons are broadly tuned to the sound cues employed in our task.

**Figure 3.**
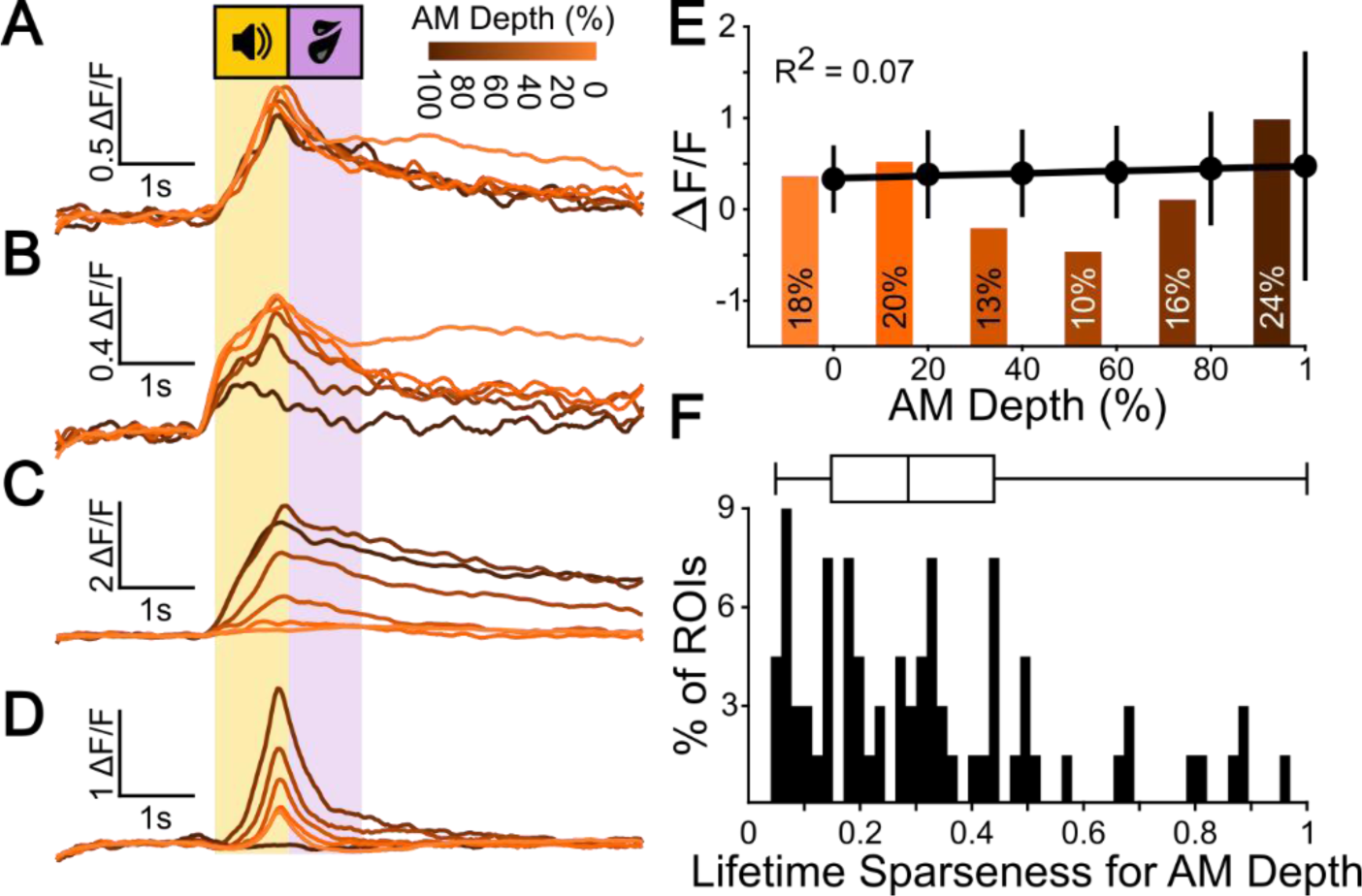
Most shell IC neurons are broadly responsive to sAM depth. A – Example ΔF/F traces for a broadly tuned representative example cell. B – The same as A for a cell tuned to high sAM depths. C – The same as A for a cell tuned to low sAM depths. D – The same as A for a cell tuned to intermediate sAM depths. E – Mean ± standard deviation of ΔF/F peak for all sound-excited cells (272) shows no linear correlation with sAM depth. Histogram bars indicate the relative proportion of significantly responsive neurons at each sAM depth. F – Lifetime sparseness for sAM depth responses of all significantly sound-responsive neurons.

### Single neuron responses are modulated by trial outcome

Many neurons showed their strongest activity modulation in the answer period of Go and NoGo trials. Neuronal activity might thus discriminate between divergent trial outcomes, such that shell IC neurons would transmit distinct signals depending on mice’s instrumental choice. We tested this hypothesis by comparing fluorescence traces averaged across trial outcomes, rather than acoustic features, for all sound and/or trial outcome responsive neurons (n = 701/909). We included all task modulated neurons in this analysis as we had no *a priori* reason to expect that trial outcome-dependent differences would be restricted to the answer period. Rather, sound-related activity might also co-vary with mice’s impending actions, in accordance with prior work demonstrating context dependent acoustic responses in IC neurons (Joshi et al., 2016; A. F. Ryan et al., 1984; Saderi et al., 2021; Shaheen et al., 2021; Slee & David, 2015).

We observed diverse trial outcome-related activity during the sound and/or answer period: Many neurons had fluorescence increases restricted to Hit and False Alarm (Figure 4A), or alternatively Miss and Correct Rejection trials (Figure 4B). Activity in these neurons thus co-varied with mice’s licking of the waterspout rather than the discriminative sound cue features, indicating that distinct shell IC populations are active depending on mice’s operant behavior. However, trial outcome related activity was not strictly yoked to mice’s actions. Indeed, other neurons had activity restricted to individual trial outcomes such as Hits (Figure 4C), or had complex activity profiles which diverged depending on whether mice’s licking action was rewarded (Figure 4D). Thus, trial outcome selectivity cannot be fully explained by a movement-related modulation of neural activity (Stringer et al., 2019; Chen & Song, 2019; Yang et al., 2020; Karadimas et al., 2020; Nelson & Mooney, 2016), but rather indicates that shell IC neurons transmit mixed representations of acoustic and higher order information related to reward, behavioral choice, motor actions, or arousal.

**Figure 4.**
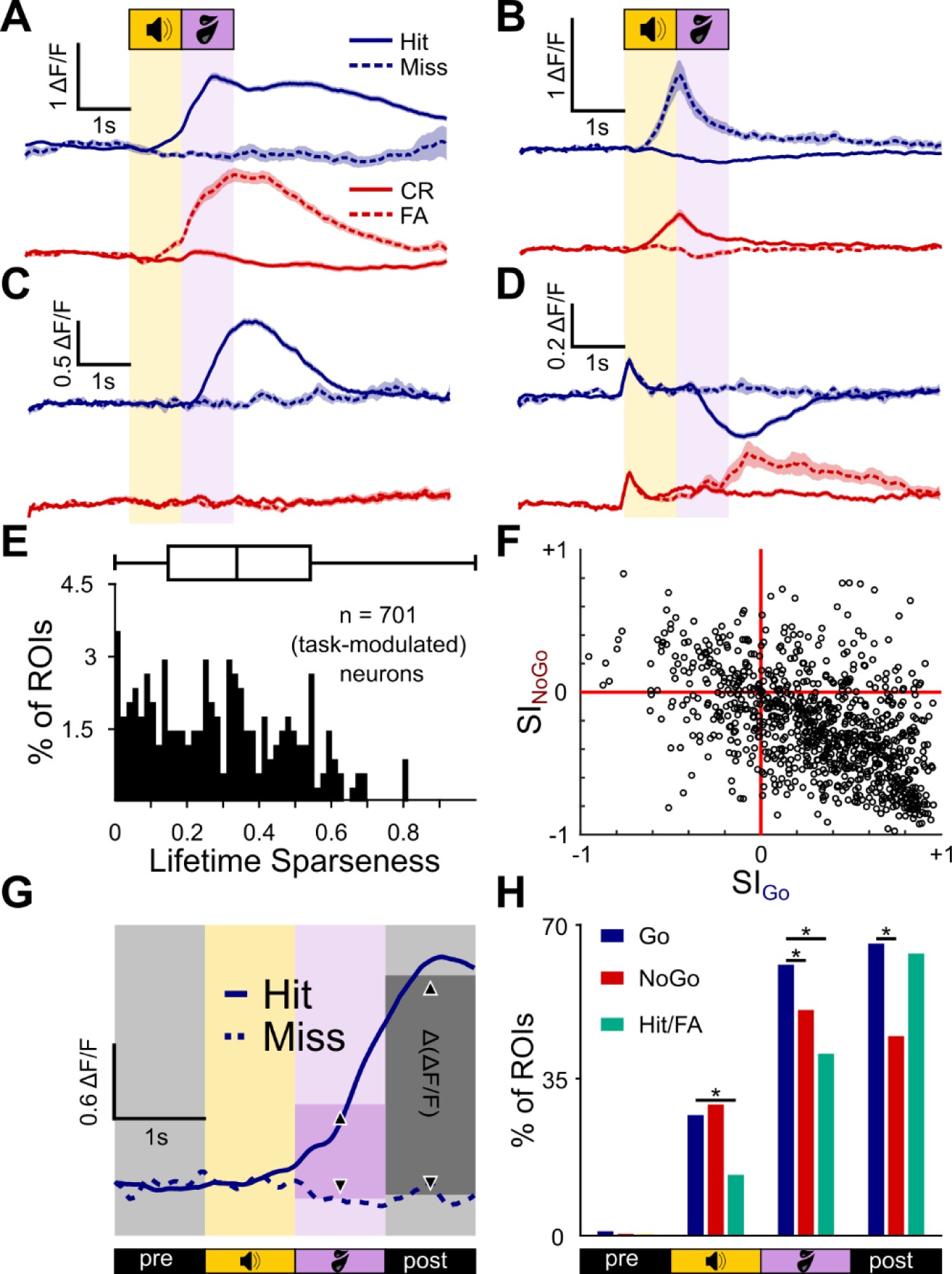
Trial Outcome selectivity of individual Shell IC neurons. A – Average ΔF/F traces of an example neuron selective for Hit and False Alarm trial outcomes. B-D) Same as A, but for a neuron responding on Misses and Correct Rejections (B), Hits only (C), or with opposing activity on Hits and False Alarms (D). E – Lifetime sparseness for outcome responses of all 701 significantly task-modulated neurons. F – Selectivity Indices on Go and NoGo trials are plotted for each neuron on X and Y axes, respectively. G – Schematic of the Δ(ΔF/F) analysis. The average ΔF/F in 1 s bins was computed on a per-outcome basis for each neuron, and compared between outcomes using a Wilcoxon rank sum test. H – The proportion of neurons with significantly different ΔF/F values for Hits and Misses (blue), Correct Rejections and False Alarms (red), and Hits and False Alarms (green) per averaging period.

Most shell IC neurons were active on multiple trial outcomes, as reflected by a low median lifetime sparseness measure in the population data (0.37; absolute derivation = 0.21; Figure 4E). We further summarized the trial outcome selectivity by measuring a separate trial outcome selectivity index (SI) value for Go and NoGo trials for each neuron. Index values range from −1 to +1 and quantify the extent to which fluorescence changes are selective for incorrect or correct trial outcomes; values of −1 and +1 indicate neurons who are only active on incorrect or correct trials, respectively. Plotting each neurons’ SI values revealed a distribution clustered towards positive and negative values on Go and NoGo trials, respectively (Figure 4F). This result indicates that correlated activity on Hits and False Alarms (as in Figure 4A) is the dominant form of trial outcome-dependent modulation, although a substantial variability in response types is clearly observable in the spread of population data.

We next quantified this diversity in trial outcome selectivity by calculating the fraction of neurons with significantly different ΔF/F values during the sound and post-sound periods of divergent trial outcomes. To this end we averaged the fluorescence values across 1 s time bins, beginning 1 s prior to sound onset and continuing until 1 s following the answer period (4 seconds total, Figure 4G). We then compared these values across Hit + Miss, and CR + FA trials (Figure 4H). In the 1 s baseline period prior to sound onset, only 1 % (7/701) of neurons showed a statistically significant difference between Hit and Miss trials; these values align with the expected false-positive rate set by the cutoff of our statistical analysis (see Methods). By contrast, 28 % (194/701) had significantly different fluorescence values during sound presentation on Hit and Miss trials, and this fraction increased to 62 % (436/701) and 67 % (470/701) during the answer- and post-answer time bins, respectively (Figure 4H, blue). Thus on Go trials, a major fraction of task-active shell IC neurons transmit signals dictated by mice’s actions rather than the features of the discriminative sound cue.

Similar results were found when comparing activity across CR and FA of NoGo trials: Although a negligible proportion of neurons showed significant differences during the pre-sound baseline period (0.4 %; 3/701), significant differences were seen in 30 % of neurons (211/701) during sound presentation (Figure 4H, red), 52 % of neurons (363/701) during the answer period, and 45 % (321/701) of neurons in the post-answer period. During sound presentation, a similar fraction of neurons showed trial outcome selectivity during Go and NoGo trials (Chi²-test, χ²(1) = 1.00, p = 0.3165). However, the fraction of neurons with differential activity in the answer- and post-answer periods of NoGo trial outcomes was significantly lower on NoGo compared to Go trials (Chi²-test, χ²(1) = 15.51, p = 0.0001 and χ²(1) = 64.40, p = 1.01*10^-15^ for answer- and post-answer periods, respectively). Thus, under our conditions, outcome selective activity preferentially occurs after sound presentation on Go trials.

We next asked whether the above differences be explained by asymmetries in mice’s licking behavior on divergent trial outcomes. If licking drives trial outcome related activity, ΔF/F responses should be similar in the answer period of Hit and FA trials where mice make similar lick actions. Consequently, very few neurons should have significant fluorescence differences during this time window. We tested this idea by comparing fluorescence signals across Hit and False Alarm trials. Pre-sound baseline differences were similar to expected false positive rates (3/701, 0.4 %), and 14 % (98/701) were significantly different during sound presentation. However, 42 % (293/701) of neurons had significant fluorescence differences during the answer period (Figure 4H, green), and these results are unlikely to be explained by differences in mice’s licking patterns during the answer period: Although mice made more licks on Hits than FA trials (7.14 ± 1.78 vs. 3.87 ± 1.80 licks/s for Hit and FA trials, respectively; p = 5.6*10^-187^, Wilcoxon Rank Sum test), most neurons remained significantly different when normalizing the fluorescence data by the total number of licks during the 1 s answer period (35 %, 247/701, Chi²-test, χ²(1) = 6.37, p = 0.186). Rather, the data indicate that many outcome selective shell IC neurons respond differently depending on whether lick actions are rewarded.

### Neural population trajectories are modulated by behavioral outcome

Our results thus far show that individual shell IC neurons transmit non-auditory and likely behaviorally relevant information, although the extent of such higher-order signals varies in magnitude across neurons. We thus asked whether task-related information is more robustly represented in population-level dynamics of shell IC activity, rather than at the single neuron level. To this end, we investigated the trajectories of neural population activity across trials. Neural trajectories are a simple way to express the network state of multi-neuronal data, and have been used in the past to compare the time-varying activity of neuronal ensembles across different experimental conditions (Briggman et al., 2005; Churchland et al., 2007; Stopfer et al., 2003). If task-related information is indeed transmitted via a population code, a network-state difference should be observable for different trial outcome conditions. We first computed a principal component analysis (PCA) of the ΔF/F traces on a timepoint-by-timepoint basis to reduce the dimensionality (Figure 5A-C). The change of the principal components over time was then defined as a neural trajectory. We generally observed a deviation of neural trajectories for Go trials (Figure 5B) depending on the trial outcome that we did not always observe for NoGo trials (Figure 5C). Surprisingly, the divergence was observed immediately following sound onset, suggesting a difference in population activity during sound presentation that varies with mice’s impending actions.

**Figure 5.**
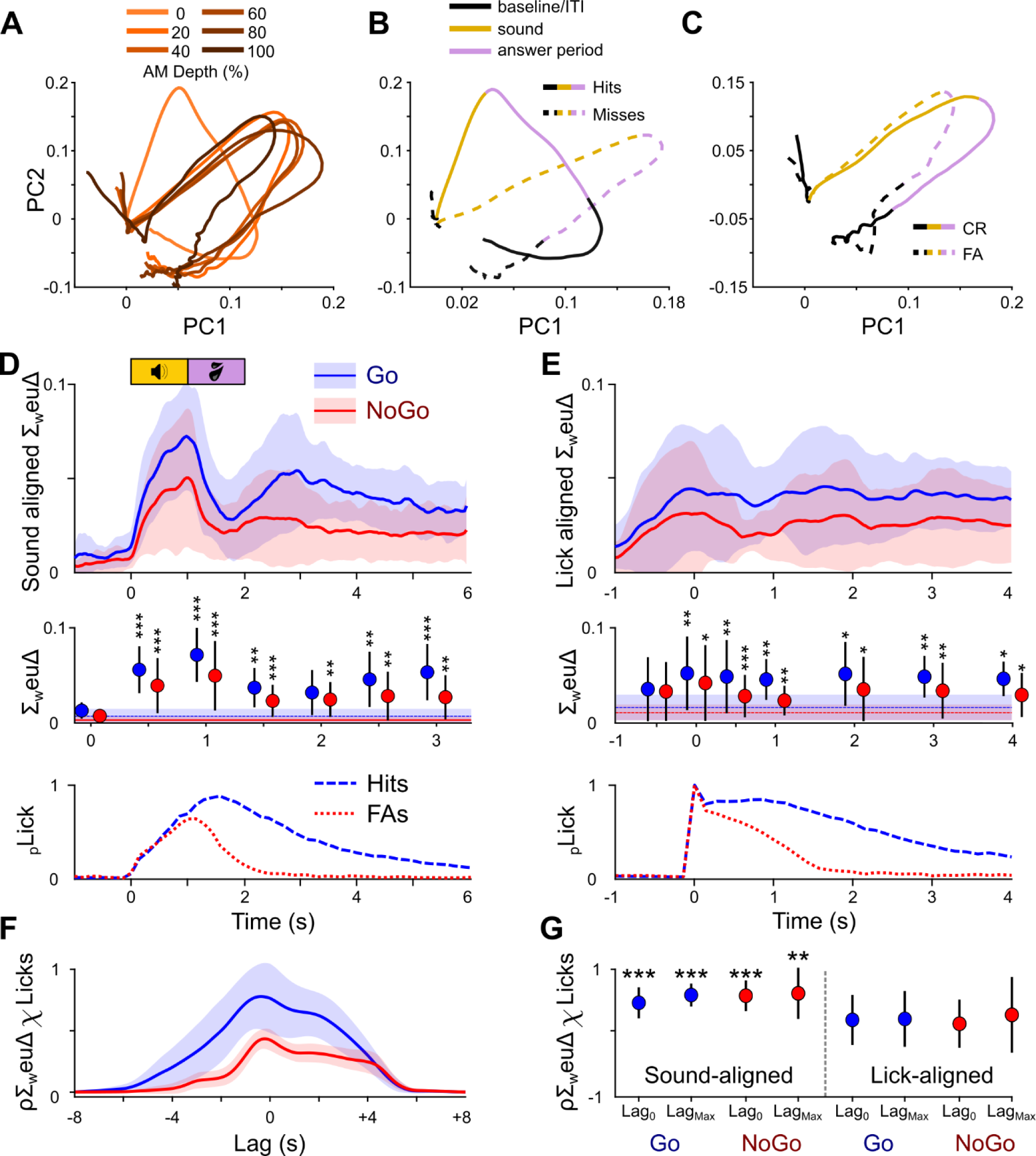
Population dynamics revealed through Principal Component Analysis show outcome-dependent differences in the processing of equal sounds. A – Example PCA-based trajectory for animal 551 sorted by AM depth. For visualization purposes, only the first 2 components are displayed, collectively explaining about 80 % of the total variance. B – The same example sorted by trial outcome for Go trials (hits/misses). Of note, the trajectories for hits and misses start to diverge immediately after the baseline. C – The same as B but for NoGo trials. D – Top: The sum of weighted Euclidean distances over all principal components over time for hits/misses (blue) and CRs/FAs (red) aligned to sound onset for all animals and sessions, plotted as mean and standard deviation. Middle: The average lick histogram on Go-(blue dashed line) and NoGo-(red dotted line) trials. Bottom: Friedman’s test followed by a Dunnett’s post-hoc test comparing the mean sum of weighted Euclidean distances against the baseline at t = −1 s. E – The same as in D, but the data were aligned to the first lick after sound onset prior to computing the PCA. F – The mean and standard deviation cross-correlation function for sound-aligned ΔF/F traces and lick histograms for Go and NoGo-trials (blue and red, respectively). G – Correlation coefficient distributions of ΔF/F traces and lick histograms for Go- and NoGo-trials for sound-aligned data (left) were significantly higher than 0 (one-sample t-test) both at 0 ms lag and at their respective maximum correlation lags, and there is no significant difference between them. In contrast, the lick-aligned ΔF/F traces and lick histograms (right) did not correlate significantly, and correlations were significantly lower than for their sound-aligned counterparts.

To quantify the trial outcome divergence in ensemble activity, we computed the Euclidean distance between the principal components of correct and incorrect trials of the same trial category: Hits vs. Misses, Correct Rejections vs. False alarms on Go and No-Go trials, respectively. We then weighted the principal components by their explained variance (_w_euΔ), and summed up the weighted Euclidean distances (Σ_w_euΔ) to compute the mean Σ_w_euΔ for Go- and NoGo trials across animals and sessions (Figure 5D). On average, we found an increasing divergence during sound presentation of both Go-(Friedman’s test, χ²(7) = 57.85, p = 4.05*10^-10^) and NoGo sounds for correct-versus incorrect trials (Friedman’s test, χ²(7) = 22.94, p = 0.0017), and a slow rejoining of the trajectories towards the end of the trial (Figure 6D, top panel). This general time course recapitulates the effect seen in the raw PCA trajectories. Interestingly, a clear second phase of divergence was also seen 2 to 2.5 s after sound onset, immediately after the answer period ended. This second phase of divergence lasted for one to two seconds before converging. Both the first and second trajectory divergences were statistically significant, as confirmed by a post-hoc Dunnett’s test comparing the baseline difference at −1 s with the peak divergences at seconds 1 and 3 (p = 1.175*10^-5^ and 5.81*10^-5^ at peak divergence 1 and 2 for Go-trials; p = 2.167*10^-6^ and 0.006 at peak divergence 1 and 2 for NoGo-trials).

**Figure 6.**
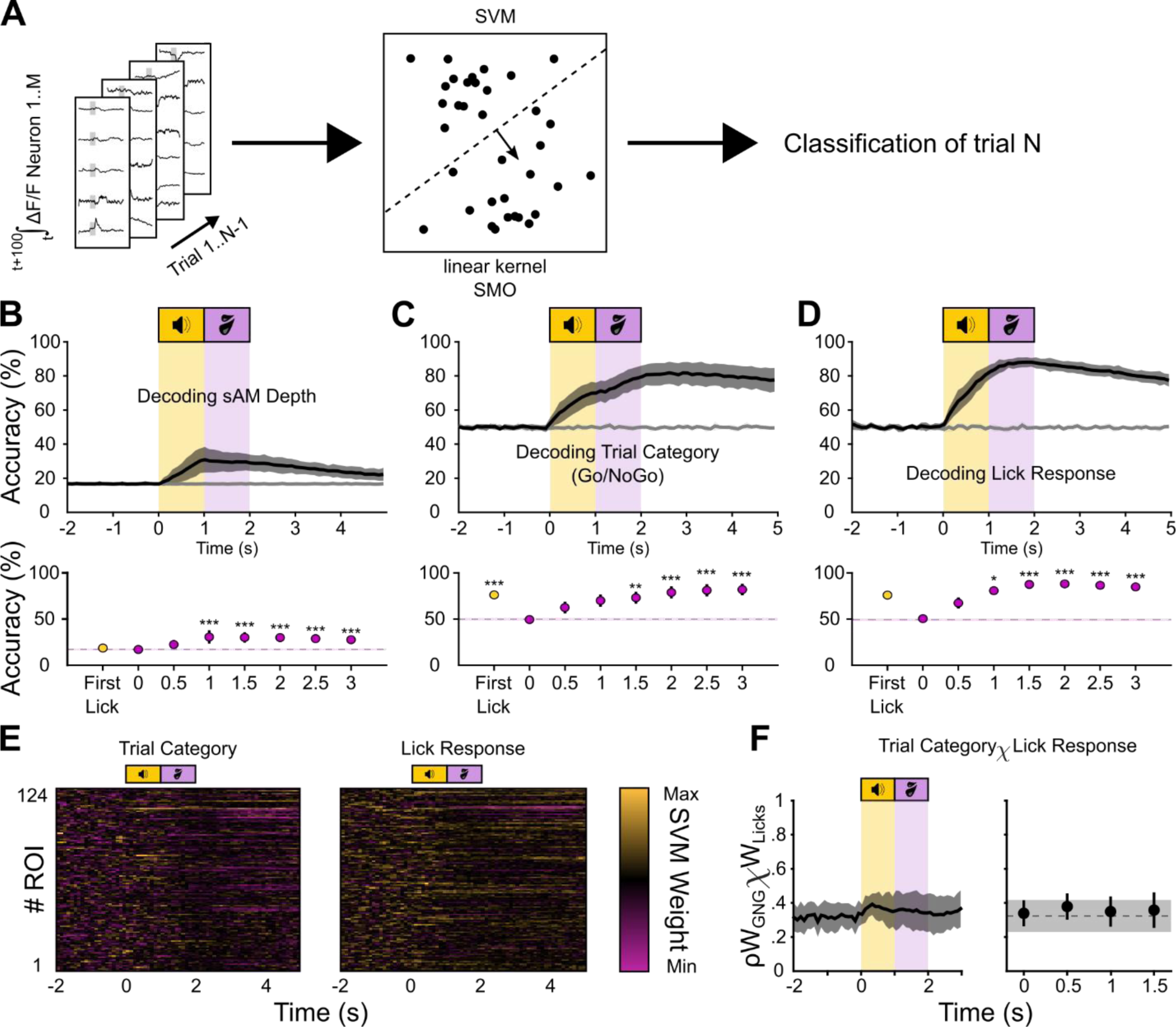
An SVM classifier can predict task-related variables from the neural activity before-, during, and after the execution of task-related behavior. A – Schematic of the SVM classifier. Training data is the integral of the ΔF/F traces of all neurons over 100 ms in a sliding window in steps of 100 ms over the trial. Accuracy is plotted over the beginning of the integration time. B – Top: Classification accuracy over time for a decoder trained to classify sAM depth. The raw accuracy was normalized to obtain the balanced accuracy (black trace), and balanced shuffled accuracy (gray trace). Bottom: Friedman-test with Dunnett’s post-hoc test comparing timepoints against the baseline accuracy at −1 s (dashed line). C, D – Same as in B, but for trial category (C) and lick response (D). E – Examples of SVM feature (ROI) weights over time for a binary classifier distinguishing Go-from NoGo-trials (left), and Lick-from No Lick-trials (right). F – Left: The mean correlation coefficients for the feature weights of the “Stimulus Category” and “Lick Response” decoders from Figure 6C and D. Each point in time represents the mean and standard deviation of the Pearson coefficients for two matched individual columns from A (feature weights at a single time point). Right: Statistics are Friedman’s test with Dunnett’s post-hoc test against baseline (t = −1 s).

A primary difference between trial outcomes is mice’s lick action. Since 440/701 (63 %) single neurons were active on hits and false alarms (Figure 4), We asked if the trajectory divergences could be explained by licking. To this end, we computed the PCA after aligning the ΔF/F traces to mice’s first lick after sound onset until the end of the answer period (or the median first-lick time of a session in trials without licks; Figure 5E). A trajectory divergence was similarly present in lick-aligned data, but divergence began before the first lick and persisted for the entire recording period for both Go-(Friedman’s test, χ²(7) = 21.64, p = 0.0029) and NoGo-trials (Friedman’s test, χ²(7) = 22.94, p = 0.0017). These results indicate that outcome selective population trajectories strikingly diverge prior to the onset of mice’s lick bouts, implying that shell IC neurons transmit different activity patterns depending on mice’s impending, rather than previously executed, actions. We next cross-correlated the averages of the lick histograms and the average Σ_w_euΔ on Go and No-Go trials to further determine the extent to which our results reflect mice’s licking patterns (Figure 5F). If the lick histogram correlates with the general curve or either of the two peaks from the sound-aligned data, we should observe one or two distinct and statistically significant maximum-correlation time points (“lags”). Indeed, we found the maximum correlation for Go-trials occurred at −0.39 s and for NoGo-trials −0.19 s, indicating that the lick histogram follows the first trajectory divergence after sound onset (Pearson’s r = 0.70, p = 8.66*10^-10^ for Go, r = 0.72, p = 2.5*10^-38^ for NoGo, Figure 5G). However, the cross-correlation function is rather broad, such that the maximum-correlation- and 0 ms lag values were not significantly different (**Figure 5G**, ANOVA, no main effect of lag time, F(1,80) = 0.99, p = 0.3225). These results suggest that the correlation might be rather unspecific, and not locked to either peak. Furthermore, when aligning the lick histogram and the ΔF/F traces to the first sound evoked lick (counted from sound onset until the end of the answer period), the correlation disappears (**Figure 5G**, ANOVA, main effect of ΔF/F alignment, F(1,80) = 16.38, p = 0.0001). This absence of correlation around the first-lick time further argues that the trial outcome-dependent, time-varying divergence of population activity does not solely reflect a motor-related component of the neural activity. Rather, the initial divergence in population trajectories (Figure 5D) may reflect a trial outcome dependent modulation of sound responses, or potentially ramping activity related to reward anticipation (Metzger et al., 2006). By contrast, the second divergence following the answer period may reflect a trial outcome-related signal that modulates IC shell neuron inter-trial activity on a timescale of seconds.

### A SVM classifier reliably decodes task relevant information from shell IC population activity

Individual shell IC neurons were often broadly responsive to sAM depth (**Figure 3**) and trial outcomes (**Figure 4**). These single neuron responses gave rise to prolonged, time-varying ensembles whose activity systematically varied with mice’s instrumental choice (**Figure 5**). Despite low individual selectivity, task-relevant information might thus still be transmitted in population activity (Robotka et al., 2023). We tested this idea by training SVM classifiers to decode specific task-relevant variables – sAM depth, trial category (Go or No-Go), and lick responses – using integrated fluorescence activity from discrete 100 ms time bins along the trial (Figure 6A). Decoding accuracy for all variables tested remained at chance level before sound onset, which is expected given that each trial’s DF/F signal was normalized to the 2 s baseline period prior to sound presentation.

We first trained the classifier to decode sAM depth, and tested if population activity transmits discriminative acoustic features at greater than chance level. The maximum classification accuracy reached was 31 ± 7 % at 1.1 seconds after sound onset (Figure 6B), thereby modestly but significantly exceeding the chance level accuracy obtained from shuffled data by 14 % (Friedman’s test, χ²(8) = 70.47, p = 3.96*10^-12^). Conversely, sAM depth could not be decoded at all when the classifier only had access to the fluorescence data from 100 ms preceding the first lick (“first-lick accuracy”), and the classifier resorts to classifying everything as BBN (Accuracy 19 ± 4 %, Figure 6B, lower panel). These results indicate that despite a rather broad sAM depth selectivity at the single neuron level, population codes might nevertheless transmit sufficient information to aid downstream circuits in discerning absolute sAM depth.

SVM classifiers also performed significantly above chance level when decoding trial category (Friedman’s test, χ²(8) = 72.58, p = 1.50*10^-12^). Interestingly, the average accuracy-over-time curve showed two separate local maxima (Figure 6B): The first plateau peaked at 70 ± 6 % at 1.1 s and likely reflected sound driven activity, as this is the earliest information available for accurate classification. The second accuracy peak rose during the answer period and reached a plateau of 82 ± 6 % at 2.7 s (31 % over chance level), suggesting that IC activity remains informative about trial category across the post-answer period. SVMs were even more robust when tasked to classify if mice licked in response to a sound (Figure 6C). Decoding accuracy peaked at 88 ± 3 % 1.9 s following sound onset (37 % above chance level, Friedman’s test, χ²(8) = 83.47, p = 9.78*10^-15^, Figure 6C), remaining elevated throughout the answer- and post-answer periods. Despite not being significant in a post-hoc Dunnett’s test against the accuracy at −1 s (p = 0.077), decoding accuracy remained high at 76 ± 3 % when using only the neural activity preceding the first lick, suggesting that the information used to decode mice’s licking may reflect preparatory motor or anticipatory activity (Metzger et al., 2006) in addition to motor-related activity itself.

Our recordings were acquired in well-trained mice who consistently performed with high Hit rates (87.1 ± 11.5%). This condition leads to a correlation between the presence of a lick response and Go trials in the training data. Thus, the lick response and trial category classifiers might achieve high accuracy via the same information, such as neural activity reflecting the acoustic features of the Go sound. In this case, licking responses might simply be predicted by proxy of their occurrence on Go trials. If true, the feature weights (= informative ROIs) assigned by the lick and trial outcome classifiers should be correlated, as classification would be based on activity in the same neurons. Alternatively, separate neurons might encode trial category and lick information, which would be reflected as a limited correlation between the feature weights of these two classifiers. We differentiated these possibilities by extracting the feature weight matrices from the lick and trial category decoders, and measuring the correlation coefficient between the “weight over ROI”-values at each time point (**Figure 6E,F**). The feature weights do not significantly correlate (**Figure 7**), indicating that lick responses are not decoded by proxy of their occurrence on go trials (or vice-versa). Rather, these results further argue that shell IC population activity transmits information related to both sound and actions.

**Figure 7.**
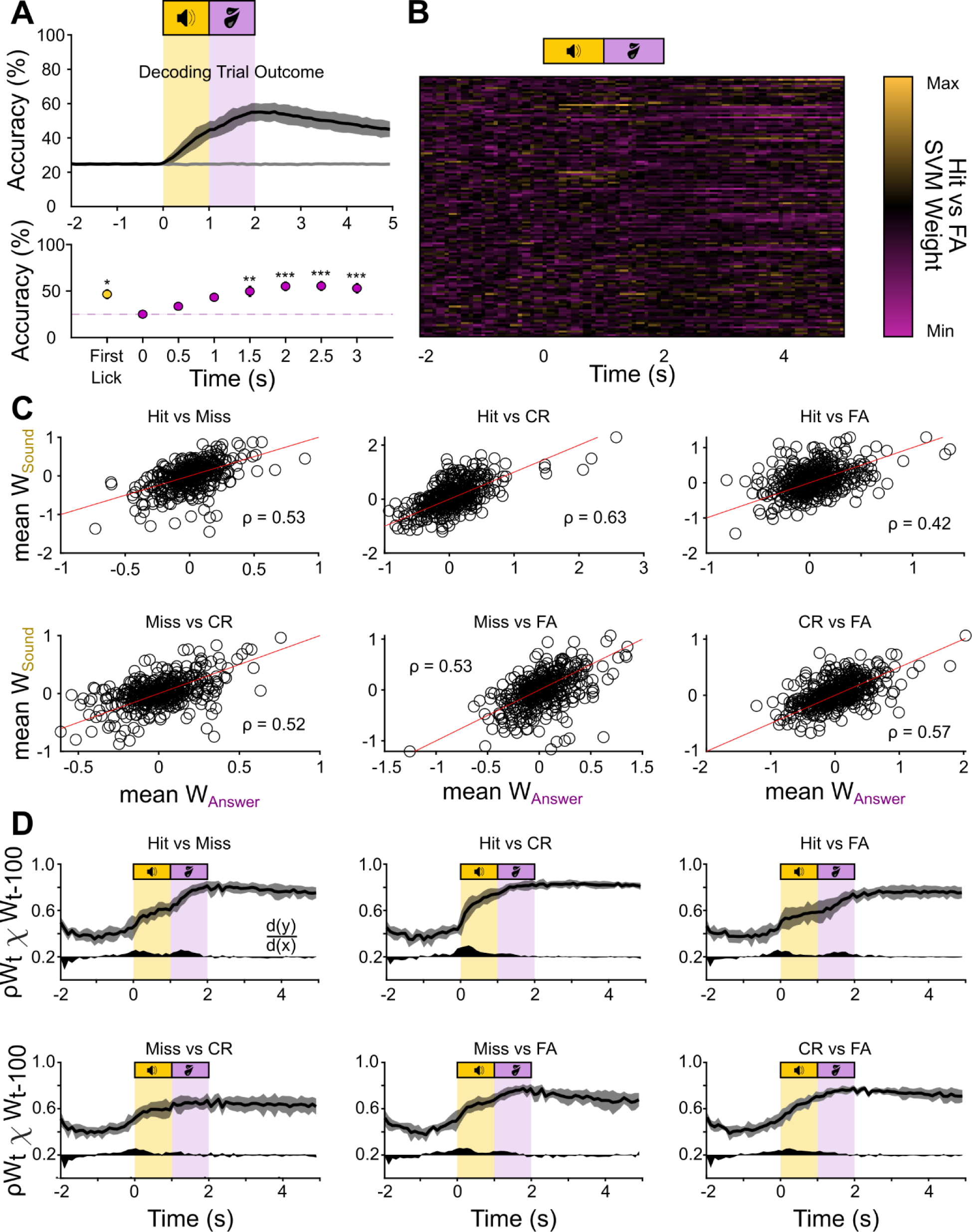
The outcome classifier uses overlapping information during the sound-and the outcome period. A – Top: Classification accuracy over time for a decoder trained to classify trial outcome. The raw accuracy was normalized to obtain the balanced accuracy (black trace), and balanced shuffled accuracy (gray trace). Bottom: Friedman-test with Dunnett’s post-hoc test comparing timepoints against the baseline accuracy at - 1 s (dashed line). B – An example set of weights for a binary classifier (Hit/FA) of the set of subclassifiers that make up the outcome-classifier. C – Mean feature weights during the sound (y-axis) and answer period (x-axis) for all ROIs for the subclassifiers distinguishing Hits and Misses, Hits and CR, Hits and FA, Misses and CR, Misses and FA, and CR and FA. Red lines are unity lines. D – Mean correlation coefficients for the feature weights of the subclassifiers at time t and time t-100 ms. Black area below the curve indicates the first derivative to visualize the steps of increased correlation in arbitrary units, with d(y)/d(x) = 0 at 0.2 on the y-axis.

### Joint population coding of task relevant signals

SVM accuracy exceeded chance levels when decoding single variables such as sAM depth, trial category, and lick occurrence (Figure 6). Thus, shell IC activity might also transmit higher order information that depends on combinations of multiple features. To test this hypothesis, we asked if a multi-class SVM could decode the trial outcome (Hit, Miss, CR, FA) from shell IC population data. In agreement, decoding accuracy for trial outcome peaked at 55 ± 5 %, 2.5 s following sound offset (Figure 7A; 30 % above chance level, Friedman’s test, χ²(8) = 82.86, p = 1.30*10^-14^). Decoding accuracy also remained above chance at 47 ± 5 % when classifier training data was restricted to 100 ms before the first lick, indicating that shell IC activity predicts trial outcomes prior to the goal-directed action. Classification accuracy of trial outcomes might reflect a uniform neuronal population whose sound responses reflect linear modulations of acoustic signals through mice’s movement, arousal, or choice. We tested this by estimating the correlation between the mean weight per neuron during the sound and answer periods, and found a significant correlation for all subclassifiers (Figure 7B, C, Pearson’s r). This correlation in the feature weight matrices suggests that the classifier uses similar populations of neurons during the sound and answer period, possibly reflecting a movement-modulated sound responsive population. However, the mean Pearson’s r was 0.53 ± 0.07, indicating a moderately large uncorrelated population. To investigate whether this uncorrelated population reflects neurons with distinctly informative responses during different times of the trial, we extracted the feature weights of the different outcome subclassifiers, and correlated the feature weights in each 100 ms time bin with those of the preceding time bin (Figure 7D). If mostly a single neuronal population drives classification accuracy, weight correlations across time should show a single increase during sound presentation that remains elevated throughout the trial. Alternatively, the observation of multiple increases in the weight correlations would indicate that partly distinct neuronal populations are maximally informative at specific trial time points. We thus plotted the weight correlation curves (Figure 7D, black curves (mean) and shading (standard deviation) and their first derivative describing the change in the curve (Figure 7D, black filled curves). Accordingly, we find two distinct steps of increased correlation, identified by local maxima in the first derivative, during the sound and answer period for most subclassifiers, suggesting that a distinct group of neurons adds trial outcome information during the answer period – likely late active neurons as those seen in Figure 4C and D.

Thus, under our conditions, shell IC population activity transmits information regarding mice’s actions in addition to acoustic signals. This activity can be used by a simple decoder to predict mice’s behavioral choice prior to their goal-directed action.

## Discussion

We have shown that in behaving mice, shell IC neuron ensembles transmit task-relevant activity that is predictive of mice’s behavioral choice, even prior to action initiation. Previous studies showed that locomotion and task engagement modulate IC neuron sound responses (Joshi et al., 2016; A. F. Ryan et al., 1984; Saderi et al., 2021; Slee & David, 2015). However, whether these effects strictly reflect an arousal- or movement-mediated change in acoustic sensitivity is unclear. We find that many neurons in the superficial IC layers are differentially active depending on mice’s behavioral choice during the response period of our task, with most task-related activity occurring following sound termination. Thus, behavioral modulation of IC neuron activity is not restricted to a scaling of acoustic responses by brain state (McGinley et al., 2015; A. Ryan & Miller, 1977; M. Zhou et al., 2014), but rather potentially reflects motor preparation, goal-directed actions, outcome evaluation, reward expectation (Metzger et al., 2006), or a combination thereof.

Approximately 40 % of shell IC neurons recorded here were not systematically responsive to the task-relevant sounds. Furthermore, sound responsive neurons were, for the most part, only weakly sensitive to increasing modulation depth of the sAM sound. These findings contrast with results in central IC neurons, where neurons are often strongly responsive to sAM sounds and steeply increase their firing rates at higher modulation depths (Krishna & Semple, 2000; P. Nelson & Carney, 2007; Rees & Møller, 1983). However, an important caveat is that our task design only tests a single modulation rate (15 Hz) of a single broadband noise carrier, and thus we were not able to establish the full range of sAM rate selectivity for our neuronal populations. Furthermore, the dynamic range of our Ca^2+^ indicators (GCaMP6f and GCaMP8s, T. Chen et al., 2013; Zhang et al., 2020) likely places an upper bound on our ability to discern spike rate differences across distinct stimuli. Despite these experimental limitations, SVM classifiers trained on shell IC fluorescence data could decode absolute sAM depth significantly above chance level. Thus, the combined activity of shell IC neuron populations may transmit signals to aid downstream circuits in reliably discriminating acoustic features. However, future studies are required to unambiguously test if shell IC neurons do or do not causally contribute to acoustic discrimination.

Neural trajectories using dimensionality reduction provide a simple measure to quantify differences in population coding that may appear subtle at the single neuron level, even for temporally overlapping stimulus features (Broome et al., 2006; Churchland et al., 2012; Stokes et al., 2013; Yu et al., 2007). Via this approach, we found that shell IC ensemble responses to physically identical sounds substantially differed depending on behavioral outcome, arguing that the sound-evoked activity of shell IC neurons is partially determined by the expected behavioral relevance. It is possible that some of this trajectory divergence reflects movement activity, preparatory or otherwise: IC neurons are known to respond to movement even in absence of sound presentation (C. Chen & Song, 2019). However, the weighted difference in population trajectories of lick- and no-lick-trials reaches a local minimum during the response window where mice’s licking response is most vigorous, thus arguing against the hypothesis that the trajectory divergences are purely motor-related. This conclusion is further supported by our observation that the first-lick-aligned neural data and lick-histograms do not correlate. Trial outcome-dependent differences in population trajectories also showed a subsequent phase of divergence occurring several seconds after mice had finished consuming the reward, *after* lick bouts had largely subsided. The mechanistic basis of this long-latency activity is unclear. However, an intriguing hypothesis is that this activity may reflect a reward- or outcome signal to update synaptic weights, or alternatively retrospective activity as reported in entorhinal cortex and hippocampus (Dotson & Yartsev, 2021; Frank et al., 2000). Alternatively, a recent theory of “adjusted net contingency for causal relations” assumes a retrospective, neutral (neither error-nor reward-based) confirmation signal (Jeong et al., 2022); A similar mechanism may explain the prolonged differences in population activity we find during correctly and incorrectly responded trials of a previously learned task. To our knowledge, our study is the first study to analyze auditory midbrain population behavioral responses in low-dimensional space. It is important to note that population trajectory differences could stem from differences in neuronal co-activity or decorrelation, which have been observed in mouse prefrontal cortex and hippocampus (El-Gaby et al., 2021; Klee et al., 2021) and are undetectable without the use of population analyses. Indeed, the general lack of specific trial outcome or sAM sound encoding in single shell IC neurons does not necessarily prohibit a neuronal population from accurately encoding complex variables (Robotka et al., 2023). These data suggest that the individually broad sound feature tuning of shell IC neurons may be advantageous for multiplexing acoustic and task-related information, such that a categorical representation of acoustic features which predict sound-driven decisions may already arise in the midbrain (Caruso et al., 2018).

What is the origin of this profuse task-relevant activity in shell IC neurons? One potential candidate is the massive system of descending auditory cortical projections. Indeed, auditory cortico-collicular neurons preferentially target the non-lemniscal shell IC layers (Bajo et al., 2007; Winer, 2005; Yudintsev et al., 2021), and are highly active during the response period in a similar instrumental task to the one employed here (Ford et al., 2022). However, auditory cortex lesions apparently reduce, but do not abolish putative non-auditory activity in cortico-recipient IC neurons (Lee et al., 2023), suggesting that shell IC neurons could inherit task-relevant activity from non-cortical sources. Accordingly, the IC receives dense projections from multiple midbrain tegmentum nuclei (Motts & Schofield, 2011; Noftz et al., 2020) which could transmit information regarding reward (Hong & Hikosaka, 2014), positive valence (Yoo et al., 2017), prediction errors (Tian et al., 2016), or behavioral outcomes (Thompson & Felsen, 2013).

Interestingly, combined responses to sound and behavioral choice, trial outcome, or unconditioned stimuli are well documented in shell IC neurons’ primary downstream targets, the non-lemniscal MGB (Barsy et al., 2020; Gilad et al., 2020; Ryugo & Weinberger, 1978; Taylor et al., 2021). This non-auditory activity is generally thought to reflect tactile or nociceptive inputs from spinal afferents (Bordi & LeDoux, 1994; Khorevin, 1980b, 1980a; Ledoux et al., 1987; Wepsic, 1966; Whitlock & Perl, 1961), and is suggested as important for associative learning and synaptic plasticity (Barsy et al., 2020; McEchron et al., 1996; Weinberger, 2011). Indeed, such non-auditory afferents could transmit a “teaching” signal to potentiate ascending IC synapses that carry acoustic information, thereby stamping in learned associations between sounds and their consequences; conceptually similar instructive signals are a hallmark of other learning related circuits (Grienberger & Magee, 2022; Raymond & Medina, 2018; Sawtell & Bell, 2008). However, our data suggest an equally plausible, alternative interpretation, namely that the auditory and non-auditory mixed selectivity found in the thalamus might partially be inherited from integrative computations upstream in shell IC neurons. Moreover, several studies suggest that IC neurons undergo plasticity during associative learning (Brainard & Knudsen, 1993; Ji & Suga, 2009; Olds et al., 1972; Vieira Lockmann et al., 2017; Vollmer et al., 2017), indicating that learning-related changes in the acoustic responses of auditory thalamus neurons (Edeline & Weinberger, 1992; Gabriel et al., 1991; Lennartz & Weinberger, 1992) could arise via synaptic plasticity in the midbrain. In tandem with our current results, these studies set the stage for future studies to test how shell IC neuron activity contributes to behaviorally relevant signals in the thalamus, and to understand the extent to which IC plasticity causally establishes a learned association between sounds and outcomes.

## Acknowledgements

The authors would like to thank Deepak Dileepkumar for technical assistance, and Drs. Michael Roberts and Luke Coddington for their helpful input on the data and manuscript. This work was supported by NIH R01DC019090, The Whitehall Foundation, and The Hearing Health Foundation.

## Author Contributions

GLQ, MMR, and PFA designed the research. GLQ, ANF, and MMR conducted the experiments. GLQ, MMR, and PFA analyzed the data. GLQ, MMR, and PFA wrote the paper.

## Notes

### Competing Interest Statement

The authors have declared no competing interest.

